# Infectious bronchitis virus regulates cellular stress granule signaling

**DOI:** 10.1101/819482

**Authors:** Matthew J. Brownsword, Nicole Doyle, Michèle Brocard, Nicolas Locker, Helena J. Maier

## Abstract

Viruses must hijack cellular translation machinery to efficiently express viral genes. In many cases, this is impeded by cellular stress responses. These stress responses swiftly relocate and repurpose translation machinery, resulting in global inhibition of translation and the aggregation of stalled 48S mRNPs into cytoplasmic foci called stress granules. This results in translational silencing of all mRNAs excluding those beneficial for the cell to resolve the specific stress. For example, expression of antiviral factors is maintained during viral infection. Here we investigated stress granule regulation by *Gammacoronavirus* infectious bronchitis virus (IBV), which causes the economically important poultry disease, infectious bronchitis. Interestingly, we found that IBV is able to inhibit multiple cellular stress granule signaling pathways whilst at the same time IBV replication also results in induction of seemingly canonical stress granules in a proportion of infected cells. Moreover, IBV infection uncouples translational repression and stress granule formation and both processes are independent of eIF2α phosphorylation. These results provide novel insights into how IBV modulates cellular translation and antiviral stress signaling.

## Introduction

During replication within a host cell, all viruses must regulate a variety of cellular processes to generate an environment that allows progeny virus to be produced to continue the infection cycle. This includes promoting pathways that are favorable to replication and overcoming intrinsic immune pathways. Cellular stress granules (SG) play an important role in regulation of gene expression by regulating mRNA translation and location as well as integrating intracellular signaling and antiviral responses and are therefore often targeted by viruses (McCormick and Khaperskyy, 2017; Walsh *et al.*, 2013). SG are cytoplasmic, non-membrane bound aggregations of mRNA associated with translation initiation factors, the 40S ribosome and RNA binding proteins. They primarily form under stress conditions that trigger the phosphorylation of translation initiation factor eIF2α (Kedersha and Anderson, 2002). There are four eIF2α kinases; protein kinase R (PKR), recognizing dsRNA, PKR-like endoplasmic reticulum kinase (PERK), sensing ER stress, heme regulated eIF2α kinase (HRI) and general control nonderepressible 2 (GCN2), activated by oxidative stress and amino acid deprivation (Deng *et al.*, 2002; Garcia *et al.*, 2007; Harding *et al.*, 1999; Lu *et al.*, 2001). Despite PKR being the assumed major kinase to activate the integrated stress response (ISR) during viral infection, PERK (Cheng *et al.*, 2005) and GCN2 (Berlanga *et al.*, 2006) have also been found to play an important role during viral infection. Phosphorylation of eIF2α prevents delivery of the initiator tRNA to initiating ribosomes, therefore inhibiting translation initiation and leading to the accumulation of stalled 48S mRNPs. SG can also be formed independently of eIF2α by interference with the RNA helicase eIF4A, which is required to unwind the mRNA untranslated region (UTR) during ribosome recruitment (Mazroui *et al.*, 2006). This can be achieved by use of the chemicals pateamine A (Low *et al.*, 2005), hippuristanol and hydrogen peroxide (Bordeleau *et al.*, 2005; Bordeleau *et al.*, 2006; Emara *et al.*, 2012). SG formation occurs in a multi-step process culminating in large compartments with dense cores held together by weak RNA-protein interactions that can merge to form SG with multiple cores. This process is driven by interactions between aggregation prone RNA binding proteins including Ras GAP SH3-domain binding protein 1 (G3BP1), T-cell restricted intracellular antigen 1 (TIA-1) and TIA-1 related protein (TIAR) (McCormick and Khaperskyy, 2017; Protter and Parker, 2016; Wheeler *et al.*, 2016). Next a liquid-like layer is formed around the core by liquid-liquid phase separation. This is achieved by interactions between RNA binding proteins containing intrinsically disordered regions (IDR) and by RNA-RNA interactions (Contu *et al.*, 2019; Lin *et al.*, 2015; Molliex *et al.*, 2015; Van Treeck *et al.*, 2018; Wheeler *et al.*, 2016). SG are highly dynamic, able to rapidly assemble, fuse and dissolve. They can act as storage sites for mRNAs allowing rapid translation reactivation upon stress resolution or mRNAs can be shuttled to sites of decay. SG are also proposed to play a role in antiviral signaling as key signaling proteins including MDA5 and PKR are known to localize to SGs and SG formation is involved in PKR activation (McCormick and Khaperskyy, 2017; Reineke *et al.*, 2015).

Viruses rely on cellular translation machinery for the synthesis of viral proteins. Therefore, the role of SG in inhibition of translation means they are often targeted by viruses to disrupt their function. Some viruses induce SG at early time points post infection but then inhibit their formation at later stages, either by inhibiting phosphorylation of eIF2α (Poblete-Duran *et al.*, 2016) or by cleaving SG scaffold proteins like G3BP1 (White *et al.*, 2007). Other viruses prevent formation of canonical SGs by redirecting SG proteins to virus driven atypical granules that co-localize with sites of viral RNA synthesis or particle assembly, benefiting virus replication (Fros *et al.*, 2012; Matthews and Frey, 2012).

Coronaviruses are positive strand RNA viruses that cause economically important diseases in humans and other species, including porcine epidemic diarrhea virus, SARS-coronavirus (CoV) and MERS-CoV. Only a few studies have been performed on the role of SG during the replication of coronaviruses. SG were found in cells infected with *Alphacoronavirus* transmissible gastroenteritis virus (TGEV). Here, viral RNA was found to be targeted to SG via an interaction with polyrimidine tract binding protein (PTB) (Sola *et al.*, 2011). SG were also found in cells infected with *Betacoronavirus* mouse hepatitis virus (MHV). Knock down of SG components, such as G3BP1 or prevention of eIF2α-phosphorylation resulted in increased viral replication, suggesting SG perform an antiviral role (Raaben *et al.*, 2007). Recently, *Betacoronavirus* MERS-CoV was found to inhibit SG formation via a process involving accessory protein 4a interaction with dsRNA and antagonism of PKR (Nakagawa *et al.*, 2018; Rabouw *et al.*, 2016).

Infectious bronchitis virus (IBV) is a *Gammacoronavirus* causing infectious bronchitis, a respiratory disease in poultry. It has been shown by others that early during IBV infection, eIF2α is phosphorylated via both PKR and PERK activation. However, at later stages, eIF2α is dephosphorylated via the upregulation of GADD153 and GADD34, promoting activity of the phosphatase PP1 (Liao *et al.*, 2013; Wang *et al.*, 2009). In addition, IBV is has been shown to shut off host translation in a process involving viral accessory protein 5b (Kint *et al.*, 2016). Despite this knowledge, the formation of SG or regulation of SG signaling during IBV replication and how this relates to regulation of translation has not been studied. Here, we present a detailed analysis of IBV regulation of cellular SG signaling and how this integrates with shut off of translation.

## Materials and Methods

### Cells, viruses and reagents

Vero cells were maintained in 1× Eagle’s modified essential medium (Sigma) supplemented with 1× L-glutamine (Gibco) and 10% fetal bovine serum (Sigma). Recombinant IBV strain BeauR has been described previously (Britton *et al.*, 2005). Inactivated IBV was generated by treatment with binary ethylenimine (BEI). Briefly, virus was incubated in 0.1 M BEI for 48 hours at 37 °C followed by inactivation of BEI by addition of 1 M sodium thiosulfate. Inactivation of virus was confirmed by RT-qPCR following infection of cells. Sodium arsenite, cycloheximide, puromycin and emetine were purchased from Sigma.

### Immunofluorescence

Vero cells seeded onto glass coverslips were mock infected or infected with IBV and incubated at 37 °C. After 1 hour, 1× BES (MEM, 0.3% tryptose phosphate broth, 0.2% bovine serum albumin, 20 mM *N*,*N*-Bis(2-hydroxyethyl)-2-aminoethanesulfonic acid (BES), 0.21% sodium bicarbonate, 2 mM L-glutamine, 250 U/mL nystatin, 100 U/mL penicillin, and 100 U/mL streptomycin) was added and cells incubated for the indicated time. Where indicated, cells were treated for 1 hour prior to fixation with 500 μM sodium arsenite or 35 μM cycloheximide or for 2 hours prior to fixation with 2 μM hydrogen peroxide. Cells were fixed in 4% paraformaldehyde in PBS, permeabilized in 0.1% triton X-100 in PBS and blocked in 0.5% bovine serum albumin (BSA) in PBS. Primary and secondary antibodies were diluted in blocking buffer. Nuclei were stained with 4’,6-diamidino-2-phenylindole (DAPI). Anti-dsRNA J2 (English and Scientific Consulting) was diluted 1:1000, anti-nsp12 (Maier *et al.*, 2013) was diluted 1:1000, anti-S2 (26.1) was diluted 1:500, anti-IBV (Abcam) was diluted 1:1000, anti-G3BP1 (BD biosciences) was diluted at 1:500, anti-G3BP1 (Sigma) was diluted 1:500, anti-eIF3η (Santa-Cruz) was diluted 1:500 and anti-eIF4G (Santa-Cruz) was diluted 1:500. Alexa Fluor-conjugated secondary antibodies (Invitrogen) were diluted 1:500. Cells were visualized using a Leica SP5 or Nikon Ti Eclipse confocal microscope. To determine the percentage cells positive for SG, cells were counted manually with at least 50 cells counted over three independent biological replicates.

### Fluorescent in situ hybridization (FISH)

Vero cells seeded onto glass coverslips were mock or IBV infected. After 24 hours, cells were fixed and labelled using the Stellaris RNA FISH simultaneous labelling protocol (Biosearch technologies). Briefly, cells were fixed in 10% formaldehyde in PBS and permeabilized in 70% ethanol at 4 °C. Cells were incubated overnight at 37 °C in a humidified chamber with hybridization buffer containing 125 nM probe and primary antibody. Cells were then washed and labelled with Alexa Fluor-conjugated secondary antibody and DAPI. Finally, cells were mounted onto glass coverslips using Vectashield and sealed with nail varnish. Cells were visualized using a Leica SP5 confocal microscope. Stellaris FISH probes with a Quasar 570 label were designed specific for the nsp15 and nsp16 region of the IBV BeauR genome.

### Cell lysis and western blot

Vero cells seeded in 6 well plates were mock or IBV infected. At the indicated time points, cells were washed once with cold PBS and lysed in 1× sample buffer (Biorad) containing DTT. Cell lysates were heated to 95 °C for 3 minutes and briefly sonicated. Proteins were separated on a 4-20 % Bis-Tris gel (Biorad) and transferred onto nitrocellulose membranes. These were blocked in 0.5% BSA or 5% milk in TBS-Tween (TBST) then incubated with primary antibody diluted in blocking buffer. Following three washes in TBS-T, membranes were incubated with HRP labelled secondary antibodies (Dako) diluted in blocking buffer. After three further washes in TBS-T, blots are incubated chemiluminescence substrate using the Clarity Western ECL Substrate (Bio-Rad). Labelled protein bands were visualized using a Vilber imaging system. Anti-IBV was diluted 1:1000, anti-eIF2α (Cell Signaling Technologies) was diluted 1:1000, anti-eIF2α-p (Cell Signalling Technologies) was diluted 1:2000 and anti-GAPDH (Invitrogen) was diluted 1:10000.

### Ribopuromycylation (RPM)

Vero cells seeded onto glass coverslips were mock or IBV infected as before. RPM was performed as described by David *et.al* (David *et al.*, 2012). Briefly, one hour prior to processing, control wells were treated with 500 μM sodium arsenite. At the indicated times post infection, cells were incubated with 18.4 μM puromycin for 30 seconds at room temperature and then incubated with 18.4 μM puromycin and 208 μM emetine at room temperature for 1 minute. Cells were washed three times with room temperature 1×BES media, fixed and processed for immunofluorescence as described above. Anti-puromycin (Sigma) was diluted 1:10000. To quantify immunofluorescence images, the puromycin signal in 100 cells was determined using ImageJ (Schneider *et al.*, 2012).

## Results

### IBV replication induces stress granules in a proportion of infected cells

Initially, the ability of IBV to induce SG during replication was assessed. Vero cells were infected with IBV and at the indicated time points cells were fixed and labelled with anti-dsRNA to detect virus infection and with anti-G3BP1 to detect SG. At each time point, infected cells were present, with the number of infected cells increasing over time, as expected (Figure 1A). In addition, at each of the time points tested, G3BP1 puncta were detected in a proportion of, but not all, infected cells with diffuse G3BP1 found in the remaining infected and uninfected cells. Subsequently, the number of infected cells with and without G3BP1 puncta was determined. The percentage of infected cells containing G3BP1 puncta was found to be between 10 and 25% (Figure 1B) and this percentage remained unchanged over the course of infection, with no statistical difference between the percentages of cells containing puncta at any time point. Therefore, IBV replication triggers the formation of G3BP1 puncta, but interestingly, only in 10-25% of infected cells.

**Figure 1.**
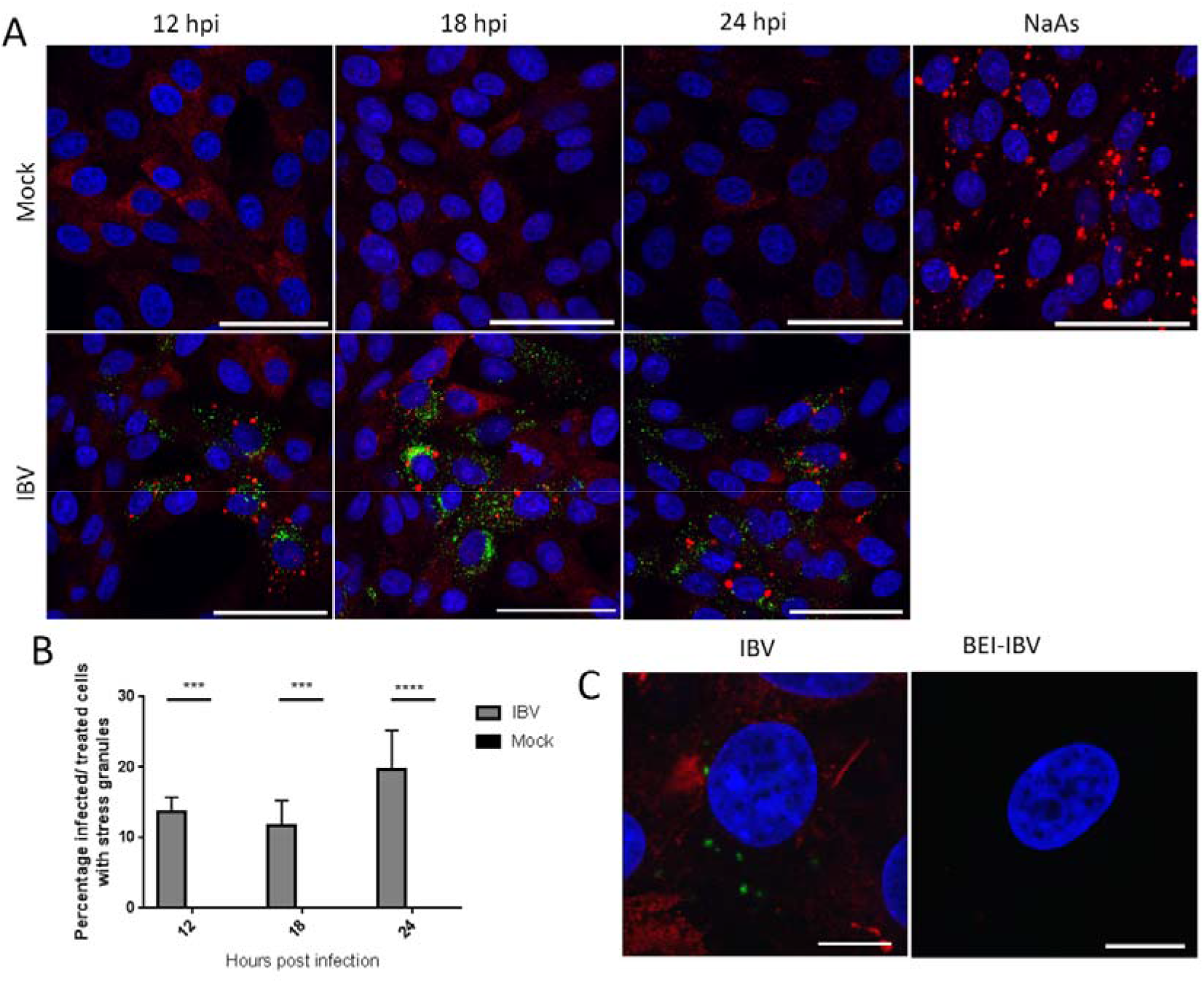
IBV infection induces stress granules in a proportion of infected cells. **(A)** Vero cells were mock infected or infected with IBV. At 12, 18 and 24 hpi, cells were labelled for stress granules (SG) with an anti-G3BP1 antibody (red) and IBV infection was detected with an anti-dsRNA antibody (green). Nuclei were stained with DAPI (blue). Positive control cells were treated with sodium arsenite (NaAs) to induce eIF2α-dependent SG. Scale bar indicates 50 μm. **(B)** Images in (A) were quantified by manual counting of SG positive cells, identified by counting infected or treated cells with G3BP1 foci. A minimum of 100 cells were counted from three independent replicates. The mean and standard deviation is shown. Asterisks indicate statistical significance as measured by one-way ANOVA, *** represents p <0.005 and **** represents p <0.0005, respectively. **(C)** Vero cells were infected with IBV or BEI-inactivated IBV. At 24 hpi, cells were labelled with anti-G3BP1 (green) and anti-IBV (red). Nuclei were stained with DAPI (blue). Scale bar indicates 10 μm.

Following identification of SG in IBV infected cells, the requirement for active virus replication in induction of granules was assessed. Cells were infected with wild type IBV or a BEI-inactivated virus. After 24 hours, cells were fixed and labelled with anti-dsRNA and anti-G3BP1. While cells infected with wild type IBV contained SG as observed before, cells infected with the inactivated virus did not (Figure 1C). Therefore, induction of SG requires actively replicating virus and is not a response by the cell to the presence of the virus particle.

### IBV replication inhibits chemical induction of stress granules

As IBV replication did not induce SG in every infected cell, it was hypothesised that IBV may be able to inhibit formation of canonical SG. To test this, cells were infected with IBV for 24 hours and prior to fixation, cells were treated with sodium arsenite for 1 hour or hydrogen peroxide for 2 hours to induce stress granule formation. Sodium arsenite induces eIF2α-dependent SG by activating the eIF2α kinase HRI. Hydrogen peroxide induces SG in an eIF2α-independent process by disrupting the eIF4F complex. Following fixation, cells were labelled with anti-dsRNA to detect virus infected cells and anti-G3BP1 to visualize SG. In uninfected cells, treatment with either sodium arsenite or hydrogen peroxide resulted in the formation of SG (Figure 2A). However, in IBV infected cells both sodium arsenite and H_2_O_2_ induction of SG was blocked with G3BP1 in infected cells remaining largely diffuse (Figure 2A). The percentage of cells containing G3BP1 foci was then determined (Figure 2B). In the absence of chemical treatment, 17% of IBV infected cells contained SG. When mock infected cells were treated with sodium arsenite or hydrogen peroxide, 83 and 74% of cells were positive for SG, respectively. However, when IBV infected cells were sodium arsenite or hydrogen peroxide treated, only 18 and 9% infected cells contained SG, respectively. Therefore, IBV infection inhibits both eIF2α-dependent and independent stress granule induction.

**Figure 2.**
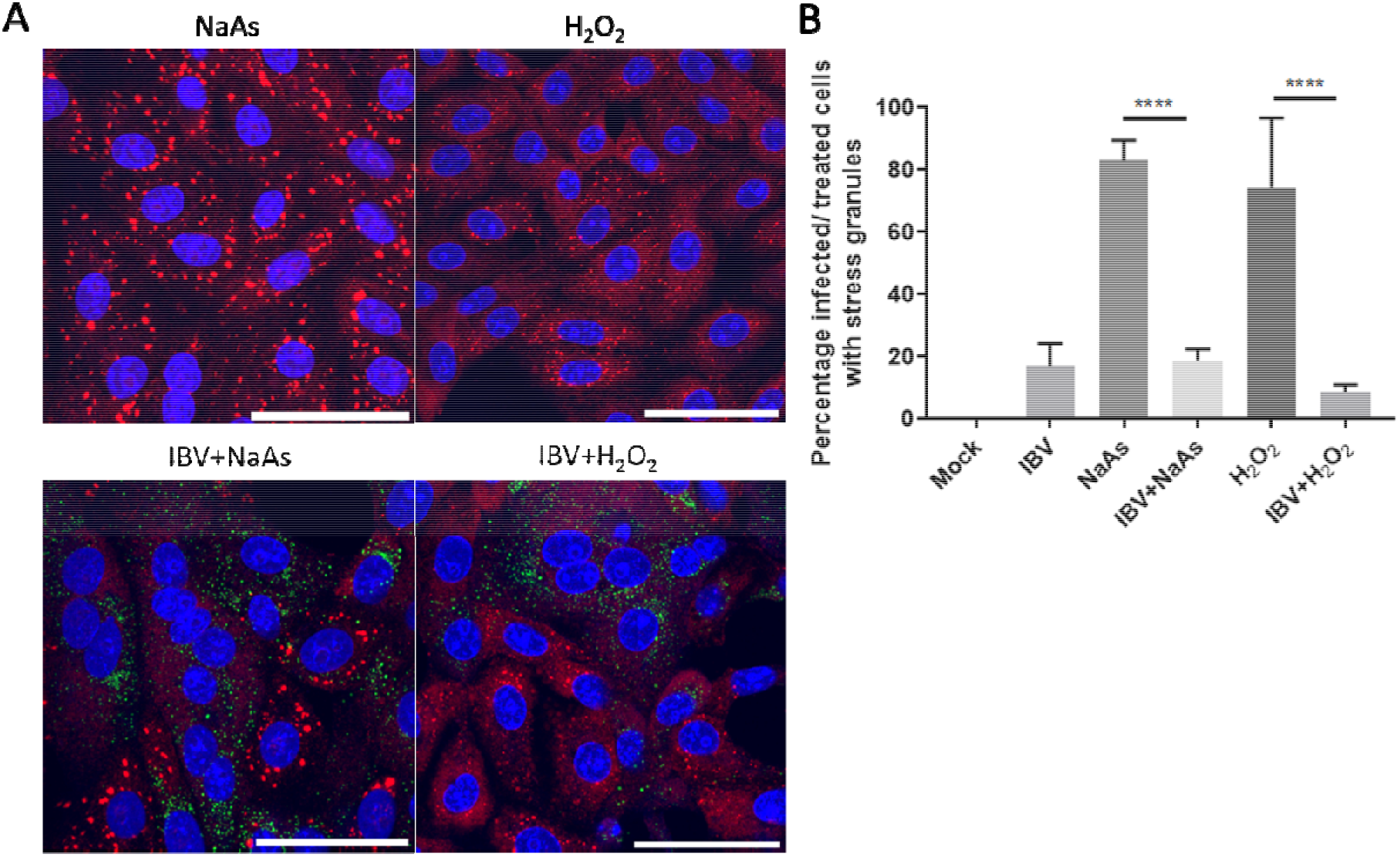
IBV inhibits eIF2α-dependent and independent stress granule induction. (**A**) Vero cells were mock infected or infected with IBV for 24 hours. Prior to fixation, cells were treated for 1 hour with 500 μM sodium arsenite (NaAs) to activate the eIF2α-dependent pathway or for 2 hours with 2 μM hydrogen peroxide (H_2_O_2_) to activate the eIF2α-independent pathway. At 24 hpi cells were fixed and stress granules (SG) labelled with an anti-G3BP1 antibody (red). IBV infection was detected with an anti-dsRNA antibody (green). Nuclei were stained with DAPI (blue) and scale bar indicates 50 μm. (**B**) Images from (A) were quantified by manual counting of SG positive cells, identified by counting infected or treated cells with G3BP1 foci. A minimum of 50 cells were counted. Data from three independent replicates. Asterisks indicate statistical significance as measured by one-way ANOVA, **** p 0.000.1

### Stress granules in IBV infected cells are canonical

Several viruses have been shown to promote the formation of specific virus-induced cytoplasmic foci by recruitment and relocalisation of many SG components including G3BP1 and G3BP2 (Fros *et al.*, 2012; Matthews and Frey, 2012; McInerney *et al.*, 2005; Poblete-Duran *et al.*, 2016). Therefore, following identification of G3BP1 puncta in some IBV infected cells, the nature of these puncta was investigated to determine whether they were canonical SG or virus-specific granules. Canonical SG contain multiple stress granule markers such as G3BP1, translation initiation factors, ribosomal subunits and mRNA. Therefore, the presence of punctate translation initiation factors eIF3η and eIF4G in infected cells was investigated. Cells were infected with IBV and after 24 hours, cells were fixed and labelled with anti-dsRNA and either anti-eIF3η or anti-eIF4G. As expected, eIF3η and eIF4G were diffuse within the cytoplasm in mock infected cells (Figure 3A). Similar to previous observations using G3BP1, in a proportion of virus infected cells, both eIF3η and eIF4G were found in cytoplasmic puncta with the remaining infected cells containing diffuse eIF3η or eIF4G (Figure 3). Therefore, IBV infection induces the formation of SG that contain multiple SG marker proteins.

**Figure 3.**
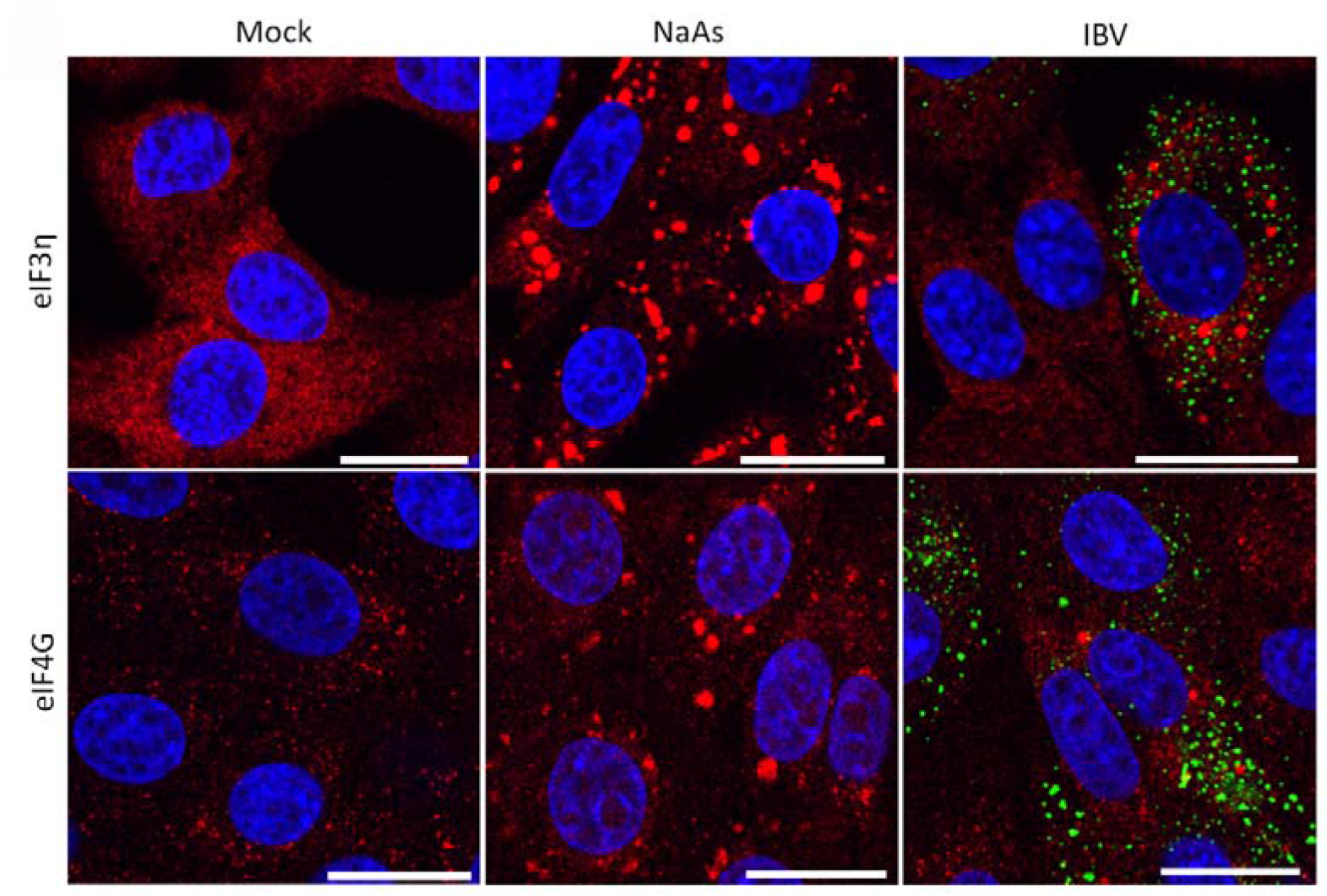
IBV induced stress granules contain multiple stress granule markers. Vero cells were mock infected or infected with IBV. One hour prior to fixation, where indicated cells were treated with sodium arsenite (NaAs). At 24 hpi, cells were fixed and labelled with dsRNA (green) and anti-eIF3η (red) or eIF4G (red), Nuclei were stained with DAPI (blue). Scale bars indicates 20 μm. Images are representative of three independent repeats.

In addition to containing multiple stress granule marker proteins, canonical SG are dissolved in the presence of cycloheximide. As mRNAs are constantly shuttled between SG and ribosomes, cycloheximide binding to the ribosome, preventing release of mRNA, inhibits recycling to SG. As a result, SG are dissolved. To further understand the nature of IBV induced SG, their susceptibility to cycloheximide treatment was determined. Cells were infected with IBV for 24 hours and one hour prior to fixation, cells were treated with cycloheximide. Cells were then labelled with anti-dsRNA and anti-G3BP1. Firstly, it was confirmed that SG induced with sodium arsenite in uninfected cells were dissolved by treatment with cycloheximide (Figure 4A). When cycloheximide treatment was applied to IBV infected cells, a significant decrease in the number of cells containing SG was observed. The percentage of infected cells containing SG was quantified (Figure 4B) and, interestingly, the number of IBV infected cycloheximide treated cells containing SG was reduced to a value not significantly different from mock cells. Together, this shows that IBV infection induces SG that contain multiple SG markers and are susceptible to cycloheximide, indicating that they are likely to be canonical SG.

**Figure 4.**
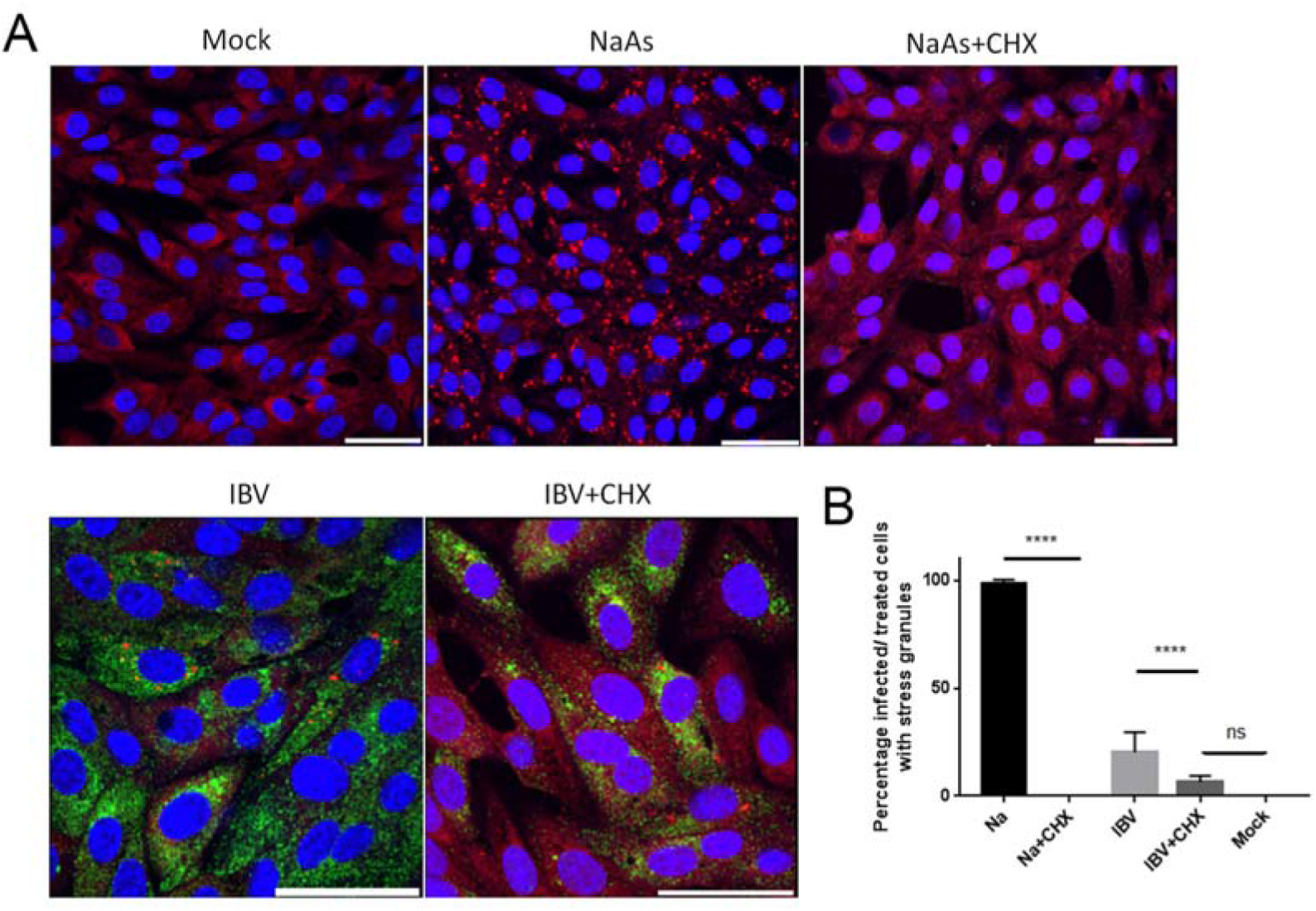
IBV induced stress granules are dissolved by cycloheximide treatment. **(A)** Vero cells were mock infected or infected with IBV. Where indicated cells were treated with sodium arsenite (NaAs). Cells were then mock treated or treated with cycloheximide (CHX) to dissolve SG. At 24 hpi, cells were fixed and labelled to detect stress granules (SG) with anti-G3BP1 (red) and IBV infected cells were detected with an anti-dsRNA antibody (green). Nuclei were stained with DAPI (blue). Scale bar indicates 20 μm. **(B)** Images from (A) were quantified to determine the percentage of cells containing SG. A minimum of 50 cells were counted. Mean and standard deviation of three independent replicates are shown. Asterisks indicate statistical significance as measured by one-way ANOVA, **** p <0.0001; ns, not significant.

### IBV genomic RNA is not diverted to stress granules during infection

It was previously found that during replication of *Alphacoronavirus* TGEV, viral RNA was targeted to virus-induced SG and this was thought to be important for their anti-viral function (Sola *et al.*, 2011). Therefore, to further understand IBV induced SG and to determine whether IBV RNA is also targeted to SG, viral genomic RNA was visualized by FISH. Cells were infected with IBV or mock infected. After 24 hours, cells were fixed and labelled with the FISH probes and anti-G3BP1 (Figure 5). Viral genomic RNA was found to be located in foci within the cytoplasm. In addition, G3BP1 puncta were detected in a percentage of infected cells, as seen before. However, no co-localization was observed between viral genomic RNA and G3BP1 containing SG. Therefore, IBV genomic RNA is not targeted to SG.

**Figure 5.**
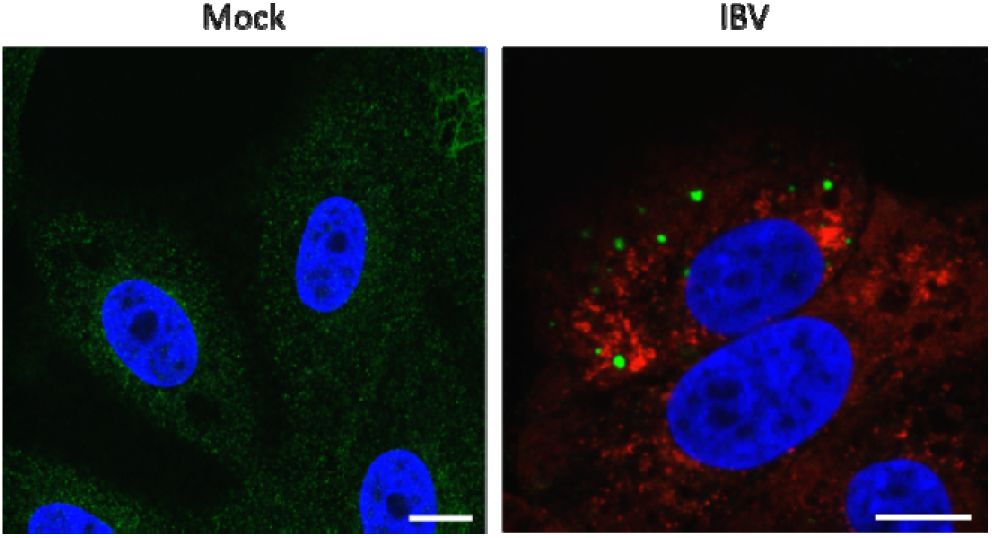
IBV genomic RNA is not diverted to stress granules during infection. Vero cells were mock infected or infected with IBV for 24 hours. IBV genomic RNA (red) was detected using FISH probes and anti-G3BP antibody was used to detect SG (green). Nuclei were stained using DAPI (blue). Scale bars indicate 10 μm. Images are representative of three independent replicates.

### Stress granules are not diverted to sites of virus replication

During the replication of several other viruses including Ebola virus, West Nile virus, dengue virus and tick-borne encephalitis virus, stress granule markers are redirected to sites of virus replication (Albornoz *et al.*, 2014; Emara and Brinton, 2007; Nelson *et al.*, 2016). To investigate whether IBV induced SG co-localize with sites of viral RNA synthesis or virion assembly, cells were infected with IBV and after 24 hours, fixed and labelled with anti-G3BP1 as well as antibodies specific for dsRNA, thought to be an intermediate in viral RNA synthesis, nsp12, the viral RNA-dependent RNA polymerase or spike protein to label for sites of progeny virus assembly. Consistent with earlier experiments, dsRNA did not co-localize with G3BP1 foci (Figure 6). In addition, G3BP1 did not to co-localize with either nsp12 or spike. Therefore, IBV does not direct SG markers to sites of virus replication.

**Figure 6.**
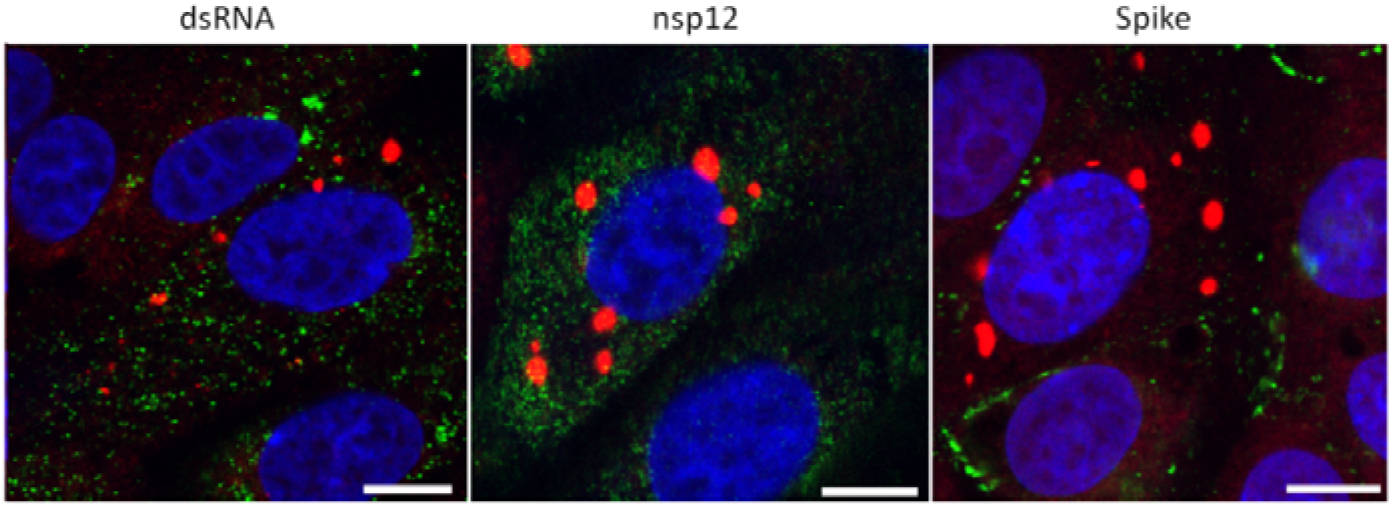
Stress granule markers are not diverted to sites of IBV replication. Vero cells were infected with IBV. At 24 hpi, cells were labelled to detect G3BP1 (red) and either dsRNA, nsp12 or spike (green). Nuclei were stained with DAPI (blue). Scale bar indicates 10 μm. Images are representative of three independent replicates.

### Stress granule formation in IBV infected cells is not correlated with translational shut-off

The formation of SG in cells is closely associated with an inhibition of translation. Previous work has demonstrated that IBV replication is associated with a shutdown of host translation from around 12 hours post infection with translation of viral proteins also ceasing by 24 hpi (Kint *et al.*, 2016). However, translational activity on a single cell level has not been characterized. Therefore ribopuromycylation (RPM) was used to visualize actively translating ribosomes over a time course of infection (David *et al.*, 2012). Cells were infected with IBV and nascent polypeptides labelled with puromycin at 12, 18 and 24 hpi followed by stalling of translating ribosomes with emetine. Cells were then fixed, stained with an anti-puromycin antibody and infected cells were detected with an anti-IBV antibody. In mock infected cells, active translation was detected with a diffuse puromycin signal throughout the cytoplasm. This signal was absent without puromycin treatment or upon treatment of cells with sodium arsenite to inhibit translation (Figure 7A). Following IBV infection at all three time points studied, two phenotypes were observed. Some cells contained the diffuse puromycin signal detected in mock infected cells. Alternatively, a proportion of infected cells showed reduced puromycin signal (Figure 7B). To enable the level of translational shut off to be determined, puromycin signal was quantified in at least 50 infected cells and surrounding non-infected cells (Figure 7C). This indicated that there was a shut off of translation at all time points. Furthermore, the degree of translational inhibition increased as infection progressed with a more pronounced shut off at 18 and 24 than at 12 hpi. However, consistent with previous work (Kint *et al.*, 2016), this single cell RPM indicates that translational shut off of infected cells is seen from 12 hpi and increases with the duration of infection. Therefore our observation that infection results in translational shut-off while only 20% of infected cells contain SGs suggests that SG assembly is uncoupled from translational arrest and/or impaired during IBV infection.

**Figure 7.**
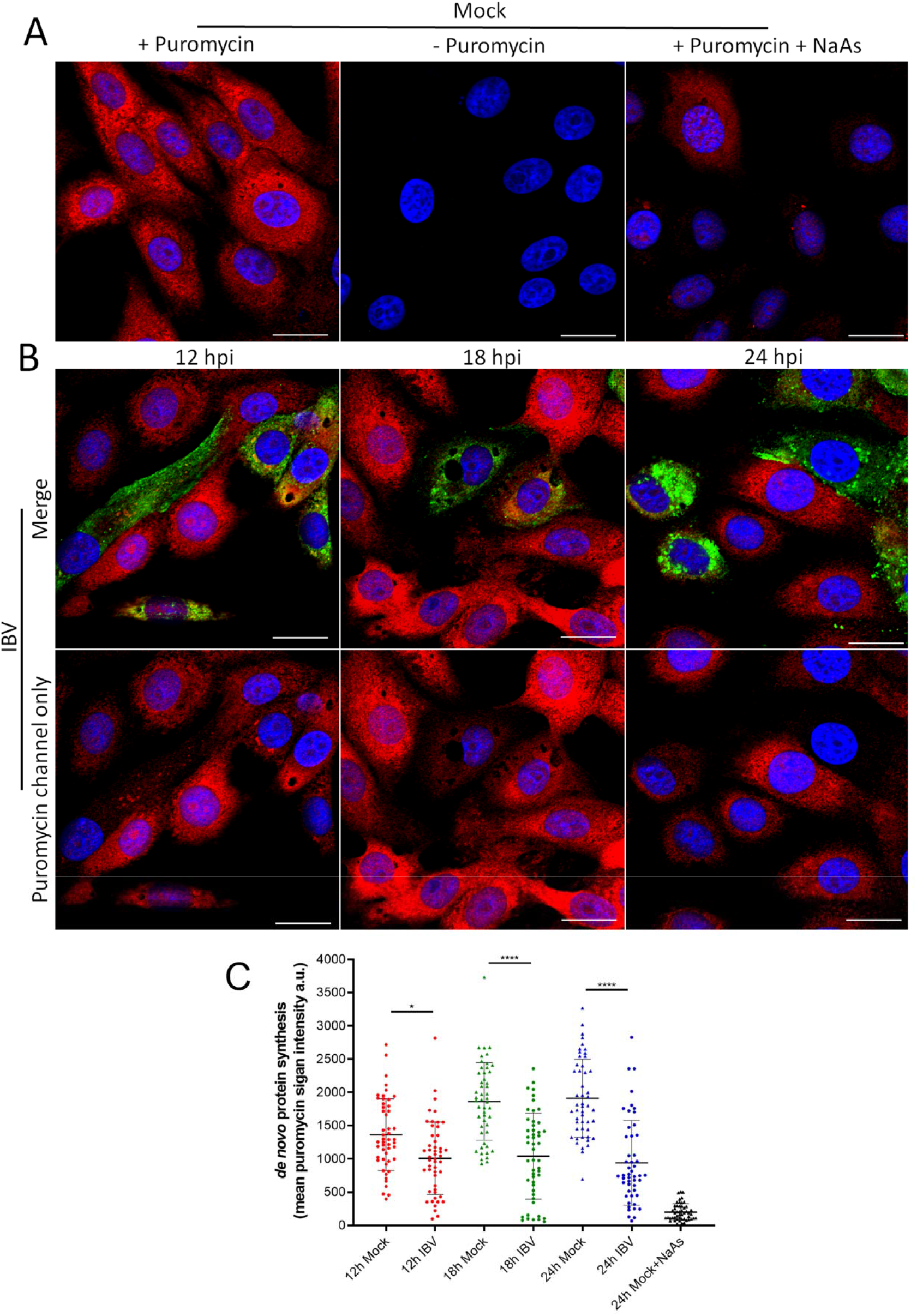
IBV infection results in translational shut-off. **(A-B)** Vero cells were mock infected **(A)** or infected with IBV **(B)**. At 23 hpi, positive control cells were treated with sodium arsenite (NaAs) to induce inhibition of translation. At 12, 18 and 24 hpi cells were treated with puromycin (or −puromycin) followed by emetine. Cells were then fixed and nascent polypeptides labelled with an anti-puromycin antibody (red) and IBV infected cells labelled with an anti-IBV antibody (green). Nuclei were stained with DAPI (blue). Scale bar indicates 20 μm. **(C)** Images from A and B were quantified. Mean puromycin channel fluorescence was analyzed in 50 cells. Data presented is representative of three independent biological replicates. Asterisks indicate statistical significance as measured by one-way ANOVA, * p =0.02; **** p <0.0001.

### Stress granule formation and translational shut-off during IBV replication are independent of eIF2α phosphorylation

Both stress granule formation and translational shut-off are usually associated with phosphorylation of eIF2α. Previous work by others has demonstrated that IBV infection results in eIF2α phosphorylation at early time points but that the virus inhibits this as infection progresses (Wang *et al.*, 2009). Therefore, the phosphorylation status of eIF2α was investigated. Vero cells were infected with IBV and lysed at 6, 12, 18 and 24 hpi. Proteins were separated by SDS-PAGE and transferred to nitrocellulose. Blots were labelled using anti-eIF2α to detect total eIF2α, anti-eIF2α-p to detect the phosphorylated eIF2α, anti-IBV to detect virus and anti-GAPDH as a loading control (Figure 8A). Total levels of eIF2α remained unchanged throughout infection. Furthermore, eIF2α phosphorylation was achieved using sodium arsenite treatment. However, during IBV infection no eIF2α phosphorylation was detected at any time point with levels remaining comparable to that of mock infected cells. Furthermore, to determine whether IBV infection actively inhibits phosphorylation of eIF2α, Vero cells were infected with IBV and then treated with sodium arsenite prior to cell lysis (Figure 8B). As before, sodium arsenite treatment of mock infected cells resulted in a significant increase in the level of phosphorylated eIF2α. When IBV infected cells were treated with sodium arsenite, there was also a significant increase in the level of phosphorylated eIF2α when compared to IBV infected untreated cells. Significantly, the level of phosphorylated eIF2α in these cells appeared comparable to that in mock infected sodium arsenite treated cells (Figure 8B). Together, this demonstrates that stress granule formation and translational shut-off observed during IBV replication both occur in the absence of detectable levels of eIF2α phosphorylation and that IBV infection does not actively inhibit eIF2α phosphorylation.

**Figure 8.**
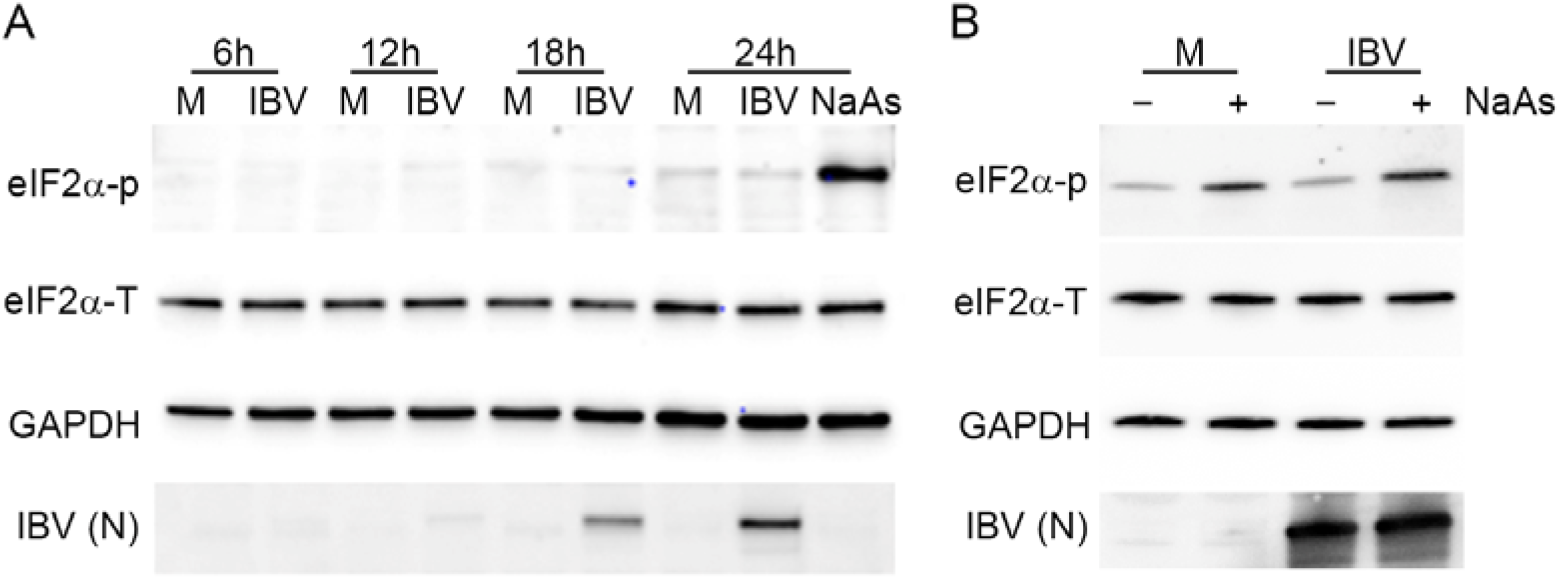
Stress granule formation and translational shut-off during IBV replication are independent of eIF2α phosphorylation. **(A)** Vero cells were mock infected or infected with IBV. At 23 hpi, cells were treated with sodium arsenite (NaAs) to induce phosphorylation of eIF2α. At 6, 12, 18 and 24 hpi, cells were washed and lysed. Samples were then separated by SDS-PAGE and transferred to nitrocellulose. Total eIF2α (eIF2α-T) was detected with anti-eIF2α (whole), phosphorylated eIF2α (eIF2α-p) was detected with anti-eIF2α-p. IBV proteins were detected using an anti-IBV antibody with a band corresponding to the IBV nucleocapsid protein shown (IBV (N)) and an anti-GAPDH antibody was used as a loading control. **(B)** Vero cells were mock infected or infected with IBV. At 23 hpi, where indicated, cells were treated with NaAs. Cells were lysed at 24 hpi and processed and labelled as in **(A)**. Blots are representative of 3 independent biological replicates.

## Discussion

Here, we present a study that furthers our understanding of how IBV regulates the important cellular pathways of the integrated stress response and translation. Firstly, we have demonstrated that IBV infection inhibits both eIF2α-dependent and independent SG formation. Several other viruses have been shown to inhibit SG signaling via regulation of the eIF2α-dependent pathway, including Kaposi’s sarcoma-associated herpesvirus, Zika virus, West Nile virus and Junin virus (Amorim *et al.*, 2017; Basu *et al.*, 2017; Linero *et al.*, 2011; Sharma *et al.*, 2017). These viruses achieve this by inhibiting activation of PKR (Sharma *et al.*, 2017) thereby preventing eIF2α phosphorylation (Linero *et al.*, 2011) or by dephosphorylating eIF2α (Amorim *et al.*, 2017). Zika virus was found to upregulate growth arrest, and DNA-damage-inducible 34 (GADD34), a component of the protein phosphatase 1 (PP1) complex, and subsequent dephosphorylation of eIF2α (Amorim *et al.*, 2017). Interestingly, in the present study, IBV was also found to inhibit eIF2α-independent SG signalling. Flaviviruses and Ebola virus also inhibit both eIF2α-dependent and independent signalling. Ebola virus achieves this via an interaction between VP35 and several SG components including; G3BP1, eIF3 and eEF2 (Le Sage *et al.*, 2017; Nelson *et al.*, 2016; Roth *et al.*, 2017). Whether any IBV proteins interact directly with SG components to inhibit SG formation remains to be determined.

Despite IBV regulation of multiple SG signalling pathways, infection results in the formation of SG in approximately 20% of infected cells. Numerous viruses are known to divert SG components to sites of virus replication to benefit the virus such as poxviruses and reoviruses (Desmet *et al.*, 2014; Katsafanas and Moss, 2007). However, although not exhaustive, our analysis suggests that the SG formed in 20% of infected cells are canonical SG produced in response to virus replication. IBV induced SG were found to contain multiple SG marker proteins and were susceptible to cycloheximide treatment, hallmarks of canonical SG. In addition, the SG markers did not co-localize with markers for viral RNA synthesis or particle assembly. Therefore, they do not appear to resemble the virus specific granules produced during the replication of other viruses. This then raises the interesting question of how SG form in this subset of cells. It is possible that viral control of SG signalling may alter over the course of infection. However, eIF2α was not phosphorylated even at early time points post infection and the SG positive subpopulation of cells remained constant at all time points tested. Therefore, this would not appear to be the case. Other viruses have also been shown previously to induce SG in only a proportion of infected cells. For example, Semliki Forest virus infection resulted in SG in 63% of infected cells at 4 hpi with a further decrease after this point (Panas *et al.*, 2012). In chronic Hepatitis C virus (HCV) infection, an oscillation of the stress response is seen in which 40% of HCV infected cells treated with interferon-α had SG but in a live time course this was shown to oscillate in a cell-specific rhythm, with 97% of these cells displaying SG at some point during infection. This appears to be a strategy by HCV to modulate the cellular stress response by PKR activated eIF2α phosphorylation or conversely, dephosphorylation of eIF2α by upregulation of GADD34 in a balancing act to prolong cell survival with oscillating stalls in translation and cell division (Ruggieri *et al.*, 2012). Therefore, it is possible that in the IBV infected cells containing SG, viral regulation of eIF2α phosphorylation or other SG signalling pathways is less efficient or these cells perhaps represent a more complex balancing act. Another possible explanation for this subset of SG displaying cells is a pre-priming of the cellular innate immunity via a paracrine signalling effect, as seen with interferon (IFN) signalling in which paracrine IFN activates the JAK/STAT pathway and upregulates interferon stimulated genes (ISGs). Analysis at the single cell level will likely be required to tease apart the mechanism of SG formation in the subset of IBV infected cells that contain them.

Other members of the coronavirus family have been found to display markedly different relationships with SG and their regulation. Transmissible gastroenteritis virus (TGEV) infection was shown to induce specific antiviral SG. In contrast to observations seen here where IBV genomic RNA did not co-localize with G3BP1, these TGEV specific granules feature an interaction between polypyrimidine tract binding protein (PTB) and viral genomic and sub-genomic RNA (Sola *et al.*, 2011). During mouse hepatitis virus (MHV) infection, translational shut-off and SG formation was also observed and MHV replication was enhanced upon infection of eIF2αS51A ^−/−^ cells or TIA^−/−^ cells in which translational inhibition and SG formation are impaired, indicating an inhibitory role for SG during MHV replication (Raaben *et al.*, 2007). Similar to our observations here, Middle East Respiratory Syndrome (MERS) CoV inhibits SG formation. This is achieved via an interaction between MERS-CoV accessory protein 4a and dsRNA, preventing PKR activation (Rabouw *et al.*, 2016). Therefore, taken together and in agreement with our findings where IBV inhibits multiple SG induction pathways, this suggests an antiviral function for SG during CoV replication.

SG usually form following translation inhibition as a result of the aggregation of stalled mRNPs, translation initiation factors and RNA binding proteins. Therefore, the translational activity of IBV infected cells was investigated. In agreement with previous work (Kint *et al.*, 2016), translation inhibition occurred during IBV replication from 12 hpi and the degree of inhibition increased as infection progressed. However, only 20% of infected cells contain SG and this remained constant across all time points studied. It is possible that some cells with reduced translational activity are the 20% observed to contain SG. However, this would not account for all the cells found to have reduced levels of translation, particularly at later time points. Instead, we consider it more likely that there is an uncoupling of SG formation and translational repression. Indeed we observe both processes in the absence of eIF2α phosphorylation indicating altered signalling for both pathways. This situation has also been observed during murine norovirus (MNV) replication where cellular translation is inhibited and canonical SG assembly is blocked through repurposing of G3BP1 independently from eIF2α phosphorylation (Brocard *et al.*, 2019; Fritzlar *et al.*, 2019). MNV translational control is achieved via the phosphorylation of eIF4E by Mnk1, which in turn is activated by the p38 kinase (Royall *et al.*, 2015; Waskiewicz *et al.*, 1997). The mechanism of IBV translational control is currently unknown. Other CoVs have been shown to inhibit translation through the action of viral non-structural protein, nsp1, which binds the 40S ribosomal subunit and cleaves host mRNA (Narayanan *et al.*, 2015). However, IBV does not express nsp1. Instead, IBV accessory protein 5b was found to be responsible for translational shut off and stability of some mRNAs tested was actually increased upon IBV infection, suggesting a completely different mechanism for control of cellular translation (Kint *et al.*, 2016).

During this project, IBV replication did not result in phosphorylation of eIF2α at any of the time points tested. Furthermore, infection was not able to limit sodium arsenite induction of eIF2α phosphorylation, showing IBV cannot actively inhibit eIF2α phosphorylation. This is in contrast to previous findings that IBV nsp2 is a weak antagonist of PKR and that GADD34 is upregulated during IBV infection resulting in decreased levels of phosphorylated eIF2α (Wang *et al.*, 2009). A subsequent study by the same laboratory found that IBV infection also induced phosphorylation of PERK, and the subsequent activation of ATF4 and the proapoptotic, GADD153, again resulting in dephosphorylation of eIF2α (Liao *et al.*, 2013). The reason for the inconsistency between our current findings and the previous work is not clear although one methodological difference in the current study is the use of sodium arsenite to induce eIF2α phosphorylation, which acts via HRI whilst the previous studies showed the activation if eIF2α phosphorylation in virus infected cells via activation of either PKR or PERK. Notably however, in our work presented here IBV replication did not induce phosphorylation of eIF2α at any time point and it is therefore not necessarily surprising that mechanisms to dephosphorylate eIF2α are also not activated. Indeed the signalling molecule required for activation of PKR, dsRNA, is known to be concealed within virus induced vesicles during coronavirus replication (Knoops *et al.*, 2008). Furthermore, interferon signalling, which also relies upon sensing of dsRNA is not activated in IBV infected cells until very late time points, consistent with shielding of dsRNA from cellular detection (Kint *et al.*, 2015). Therefore, the activation of these various cellular signalling pathways in response to IBV infection is likely to be prevented, consistent with our findings.

## Conclusions

In the present study we have demonstrated that IBV replication effectively blocks both eIF2α-dependent and eIF2α-independent SG signalling pathways. In addition, IBV replication results in a shut-off of translation. However, interestingly in a proportion of infected cells, canonical SG are formed that do not localize with sites of viral replication and do not contain viral RNA. This raises the interesting future possibility of being able to study the composition and function of canonical cellular SG in virus infected cells. In addition to these findings, in IBV infected cells, both translational repression and SG formation were found to occur in the absence of eIF2α phosphorylation, although IBV replication was not able to actively inhibit eIF2α phosphorylation. Therefore, IBV infection of cells results in a dysregulation and uncoupling of several important cellular signalling pathways. The mechanism behind this dysregulation remains to be determined but we have furthered our understanding of how IBV changes the cellular environment to make it favorable for virus replication.

## Author Contributions

conceptualization, H.J.M. and N.L.; methodology, M.J.B., N.D., M.B., N.L. and H.J.M..; validation, M.J.B., N.D., N.L. and H.J.M; formal analysis, M.J.B.; investigation, M.J.B., N.D., N.L. and H.J.M.; resources, H.J.M. and N.L.; writing—original draft preparation, M.J.B. and H.J.M.; writing—review and editing, M.J.B., N.D., M.B., N.L. and H.J.M.; visualization, M.J.B and H.J.M.; supervision, H.J.M. and N.L.; project administration, H.J.M.; funding acquisition, H.J.M. and N.L.

## Acknowledgments

The authors would like to thank Ambi Batra for providing BEI-inactivated IBV.

